# Gene family evolution reveals dietary adaptations in butterflies and moths

**DOI:** 10.1101/2025.01.15.632883

**Authors:** Yi-Ming Weng, Jose I. Martinez, Amanda Markee, David Plotkin, Yash Sondhi, Andrew J. Mongue, Paul B. Frandsen, Akito Y. Kawahara

## Abstract

Butterflies and moths (Lepidoptera) are a hyperdiverse lineage of nearly 160,000 described species and their evolutionary success is postulated to be tightly correlated with the radiation of their primary host — angiosperms. Previous studies found that a significant number of emergent gene families are specific to Lepidoptera, with many genes linked to odorant receptors and peptidases, suggesting that such genetic innovations may be linked to their diversification. Here, we use genomic resources to identify lineage-specific gene families in four nested lepidopteran clades (e.g. Lepidoptera, Glossata, Ditrysia, and Apoditrysia). Among nearly a hundred gene families specific to each group, a handful of gene families have specific and interpretable functions. We found that many digestion-related gene families emerged early in the evolution of Lepidoptera, followed by genes associated with detoxification and immunity. This result aligns with the evolutionary transition in ancient Lepidoptera diets to terrestrial plants, highlighting the emergence of detoxification mechanisms in the megadiverse Ditrysia as a critical adaptation driven by the proliferation of plant chemical defenses. We also found gene families originating from horizontal gene transfer (HGT) events, likely introduced from bacteria and fungi to the common ancestors of Lepidoptera and Ditrysia, respectively. These HGT-derived genes likely played a pivotal role in supporting the dietary transition from algae, diatoms, and aquatic plant debris to fungi and primitive terrestrial plants, ultimately enabling the adaptation to the most dominant angiosperm species.

## Introduction

Butterflies and moths (Lepidoptera) constitute the second largest order of insects with nearly 160,000 described species in 43 superfamilies and 133 families (1). Lepidoptera is the most dominant herbivorous insect order, with nearly all species feeding on plants during their larval stage, and adults relying heavily on nectar and other plant-derived nutrients. Lepidoptera is believed to have diverged from a common ancestor shared with caddisflies (Trichoptera) during the Late Carboniferous period, with an estimated crown age of approximately 300 million years (2, 3). The chewing mouthparts of the earliest diverging “mandibulate” moths, shared with caddisflies, are considered the ancestral condition in the order, and this trait is retained only in the three oldest extant families: Micropterigidae, Agathiphagidae, and Heterobathmiidae, which are primarily fungivores or detritivores (4). Subsequently, the overwhelmingly dominant siphoning mouthpart, or proboscis, evolved prior to the division of Glossata, around 240 Ma. Within Glossata, Ditrysia constitute 98% of extant ordinal species diversity, and nearly all species in Ditrysia rely on angiosperms during their larval and adult stages. The dependence on angiosperms has led researchers to hypothesize that a key evolutionary event enabled ancestral Lepidoptera to transition to angiosperm feeding. However, the precise mechanisms underlying this shift remain unknown.

Genomes of extant species may shed light on how the ancestral lepidopteran may have shifted to angiosperm feeding. A recent study revealed a remarkable number of unique emergent gene families only found in Lepidoptera, surpassing the number found in insect orders with similar or greater species diversity (e.g. Coleoptera, Diptera, Hymenoptera) (12). These emergent gene families were found to be associated with peptidases and odorant binding hypothesized to be related to protein digestion and host detection, implying their role in host plant adaptation. However, the study only included five ditrysian species, and none of the non-ditrysian lineages were represented, thereby limiting the ability to test whether these genes are unique to Ditrysia. Key questions remain, such as when these gene families first emerged and diversified, and how they relate to host-plant associated diversification in the order. By comparing genomes of extant species from both early and recently derived lineages, we can understand how these emergent gene families correlate with the evolutionary history of Lepidoptera.

In this study, we compiled 431 genomes of Amphiesmenoptera (Lepidoptera and its sister-group, Trichoptera), to investigate gene family evolution and its relationship to major evolutionary events in Lepidoptera. We test the hypothesis that the remarkable diversification of Ditrysia is linked to adaptations for feeding on angiosperm host plants. We identify lineage-specific gene families associated with key evolutionary transitions in Amphiesmenoptera, Lepidoptera, Glossata, Ditrysia, and Apoditrysia. We also found lineage-specific gene families derived from horizontal gene transfer (HGT), likely from bacteria and fungi. These genes play roles in food digestion, embryo development, and mating behavior (13, 14). We found that digestion-related gene families emerged early in Lepidoptera evolution, followed by detoxification and immunity genes in derived Ditrysia, highlighting the adaptation to plant-based diets. By analyzing both conserved and rapidly evolving gene families, we reveal significant genetic innovations that underpin major evolutionary steps closely tied to the radiation of angiosperms.

## Results

### BUSCO score assessments and Lepidoptera phylogeny

In order to understand the evolution of Lepidoptera genome evolution, we first obtained genome assemblies of 431 Lepidoptera and Trichoptera species. We found that most had high BUSCO completeness when using the endopterygota_odb10 database (**Table S1**). We also used the lepidoptera_odb10 database, which resulted in Trichoptera having lower BUSCO scores, as expected, given that the gene set was built based on Lepidoptera specific single copy orthologs. Interestingly, we observed a gradual decrease in BUSCO completeness from Apoditrysia to the most ancestral mandibulate moth lineage (Micropterigoidea), which exhibited BUSCO scores similar to Trichoptera (**Figure 1**). This is likely because those early derived lineages including Micropterigoidea were not used in the generation of the ortholog gene database. We determined representative species by choosing the species with the highest BUSCO scores (endopterygota_odb10) in each family. As a result, 62 species were selected to represent 48 families across 27 superfamilies (including 21 Lepidoptera superfamilies) for phylogeny reconstruction (**Table S1**). The species tree topology based on the concatenated supermatrix of 1,439 BUSCO single copy genes (endopterygota_odb10) was congruent with recently published phylogeny built with transcriptomic data (3), with high support on almost all branches (**Figure 2**). The crown ages of Lepidoptera, Glossata, Ditrysia, and Apoditrysia were 276, 233, 157, and 123 million years ago (Ma), respectively.

**Figure 1.**
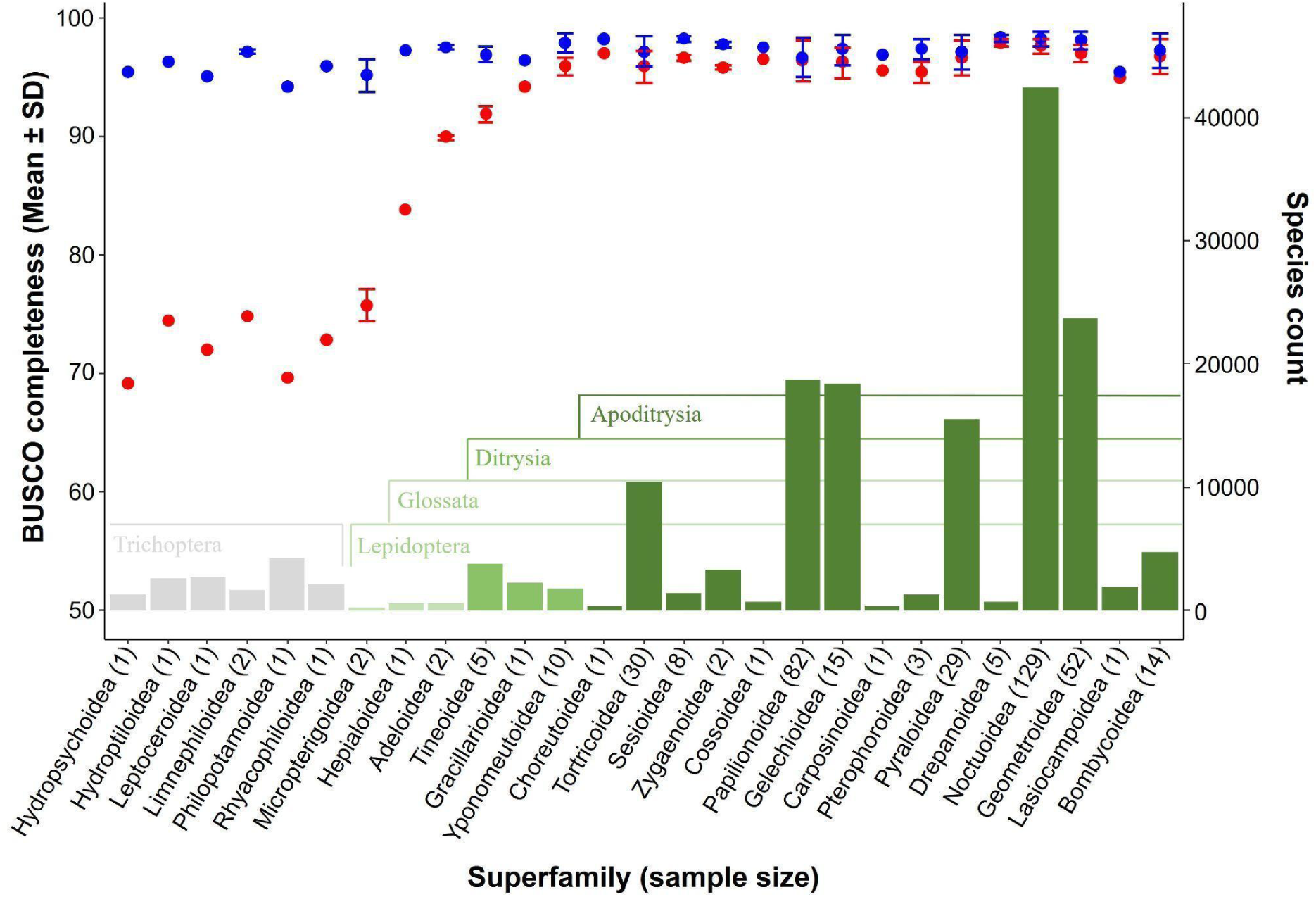
Comparisons of BUSCO complete scores between results from lepidoptera_odb10 (red dots) and endopterygota_odb10 (blue dots) databases. BUSCO complete scores were averaged by superfamily with the standard deviation represented by the error bars, while sample size is marked on the superfamily names on the X axis. Color bars denote major lineages including Trichoptera (gray), and four key, nested lepidopteran groups (Lepidoptera, Glossata, Ditrysia, and Apoditrysia). The height of each bar shows the estimated species diversity in each superfamily based on references (2, 44).

**Figure 2.**
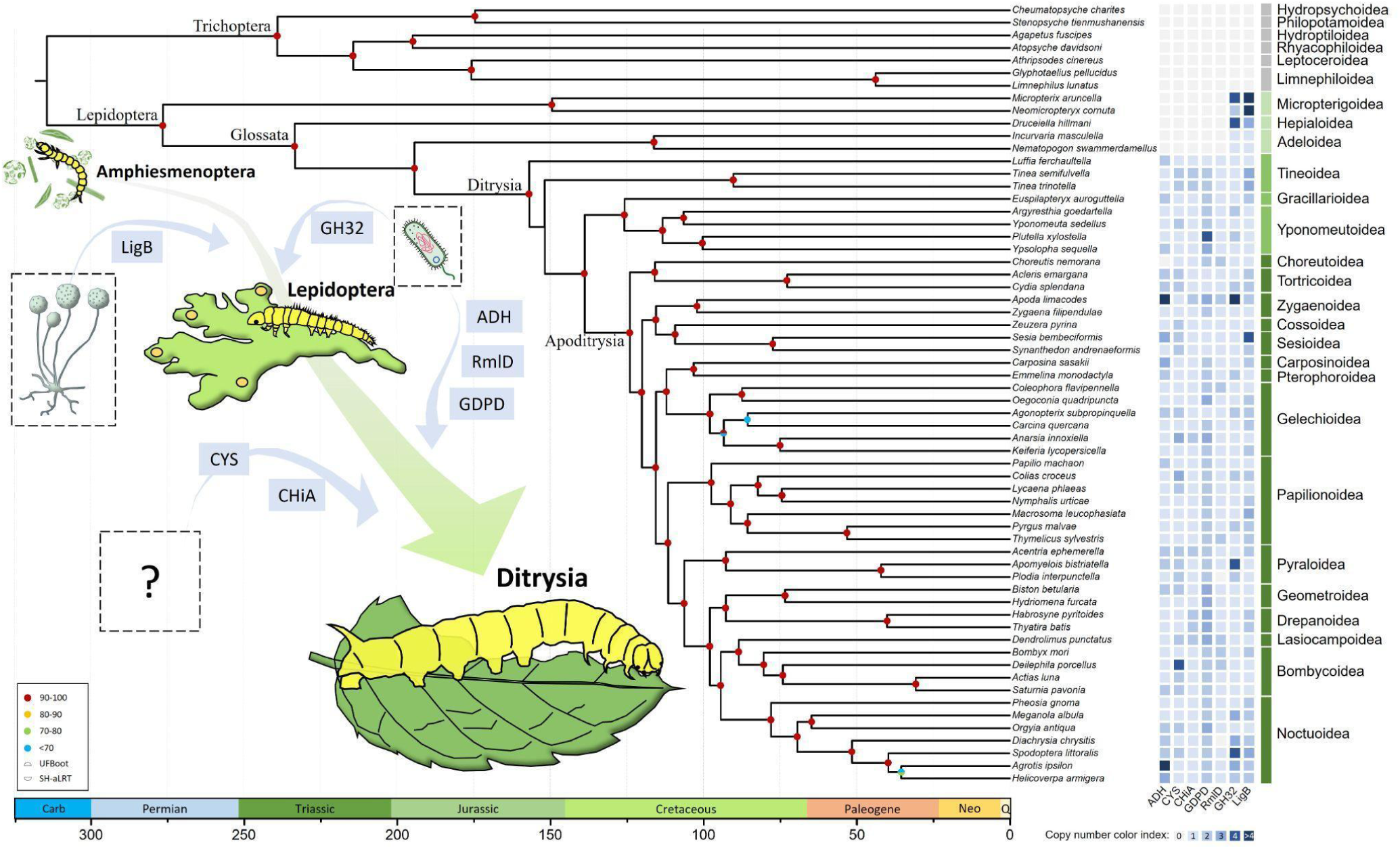
A time-calibrated species tree based on 1,439 single copy BUSCO genes. Timing of the horizontal gene transfer (HGT) events from bacteria and fungi are indicated. This tree also illustrates the general transition of their diet from algae, diatoms, or aquatic plant debris (Amphiesmenoptera) to primitive terrestrial plants (Lepidoptera), and later to angiosperms (Ditrysia). The heatmap on the right shows gene copy numbers of HGT-derived genes. Illustration of the caterpillar was adopted from reference (69).

### Gene family inference from re-annotated genomes

To standardize protein coding gene annotation, we re-annotated the softmasked genomes of the 62 representative species using the BRAKER3 pipeline. BUSCO scores of the predicted protein coding gene models mirrored the BUSCO scores of their respective genome assembly, where “lower” Lepidoptera and Trichoptera were less BUSCO complete when using the lepidoptera_odb10 database. In general, gene models predicted through the BRAKER ETP pipeline had higher BUSCO completeness than those of BRAKER EP pipeline, with marginal differences. Orthologous inference from these 62 species’ gene models included 50,523 Phylogenetic Hierarchical Orthogroups (HOGs), where 19,265 HOGs comprised more than 10 sequences.

### Rapidly evolving gene families

To investigate gene family evolution in Lepidoptera, we identified rapidly evolving gene families. We characterized HOGs with significant changes in repertory size (gene copy number). Gene families with significantly rapid repertory size changes at all major lineages were mostly associated with retrovirus activity, including *Reverse transcriptase*, *Retrotransposable element*, *Ribonuclease H protein*, and *Pao retrotransposon peptidase* (**Table S2**). In Trichoptera and lower lepidopteran lineages, some gene families associated with nutrition management, immune responses, and detoxification evolved rapidly. For example, *C-type lectin*, *UDP-glucoronosyl*, *UDP-glucosyl transferase*, and *Insect cuticle protein* were significantly contracted in Trichoptera, while *Enoyl-reductase*, *Trypsin-like serine protease*, and *Insect cuticle protein* expanded their repertory size rapidly during the time when the proboscis evolved in the early Lepidoptera (i.e., Glossata). In the hyperdiverse lineage, Ditrysia, chemosensory genes such as the *7tm Odorant receptor* and *Gustatory receptor* were significantly expanded. In Apoditrysia, *Peptidase S1 family*, *Immune response gene C-type lectin*, and Putative *detoxification gene cholinesterase-like* are expanded, while the Putative *peptidase DUF1758* was contracted.

### Newly emerging, lineage-specific gene families

For lineage-specific gene families, defined by their uniqueness and prevalence within the lineage (i.e., HOGs that contain genes present in >95% of species in the focal group and 0% in the other species), 82-120 HOGs were specific to each major lineage (57-89 with EggNOG function), except for the youngest lineage, Apoditrysia, which had only five unique HOGs (four with EggNOG function) (**Table S3**). Many of the annotated gene ontology (GO) functions (biological process) from these lineage-specific genes are linked to a diverse range of biological processes, encompassing a broad spectrum of functions. Nevertheless, there are many terms that show specific functions with a direct implication to ecological adaptations such as genes involved in chemosensory, digestion, detoxification, response to stress, immune related, and cuticle properties (**Table S4**). Among the Trichoptera-specific gene families, several are associated with cuticle formation, including *Cuticle protein 18.7-like*, *Larval cuticle protein A2B-like*, *uncharacterized protein GBIM_19595* and possibly *Choline transporter-like protein 1*. Other genes families specific to Trichoptera include cold-stress related genes such as *Facilitated trehalose transporter Tret1-like* and *Endoglucanase E-4-like*; detoxification related genes such as *UDP-glucuronosyltransferase 2B15-like*, *cytochrome P450 4C1-like*, and *Venom allergen 5-like*; immune related genes such as *Serine protease inhibitor dipetalogastin-like* and *Serpin*; digestion related genes such as *Myrosinase 1-like*; and sensory genes such as *Odorant receptor 19a-like*.

In Lepidoptera, many *cytochrome P450* HOGs are found to be unique, with one annotated with GO term “response to camptothecin (an alkaloid)” and others linked to responses to various xenobiotic stimuli. Several other unique gene families are potentially associated with cuticle properties, including an unnamed gene containing a chitin-binding domain, *Endocuticle structural glycoprotein SgAbd-3-like* and *Pyrimidodiazepine synthase-like*. HOGs specific to Lepidoptera also include chemosensory genes such as *Chemosensory protein* (*Insect pheromone-binding family, A10/OS-D*) and an unnamed gene in the carboxylesterase family which is involved in carboxylic ester hydrolase activity, as well as digestion related genes including *membrane alanyl aminopeptidase-like* and *serine/threonine-protein kinase unc-51*. Remarkably, two HGT-derived genes, *Catalytic LigB subunit of aromatic ring-opening dioxygenase* (*LigB*) and *Glycosyl hydrolases family 32* (*GH32*), are specific to Lepidoptera and related to plant tissue digestion, with novel EggNOG functions absent in Trichoptera (**Figure 2, Figure S1-S2**).

In Glossata, many newly formed gene families are involved in cuticle formation and pigmentation, such as *Peroxisomal N*(*1*)*-acetyl-spermine/spermidine oxidase-like*, *Endocuticle structural glycoprotein ABD-4-like* and *ABD-5-like*, an unnamed cuticle protein, and an unnamed protein with chitin-binding domain type 2. Other notable genes specific to Glossata include vision genes such as *Opsin-1* (long wavelength opsin) and *Disco-interacting protein 2*; chemosensory related genes such as *Pickpocket protein 28-like*, *Homeotic protein empty spiracles-like*, *Glutamate receptor 1-like*, and *Odorant receptor*. For host plant related genes, we identified an unnamed protein with peptidase inhibitor activity, *Trypsin-like serine protease*, and *Myrosinase 1* (likely associated with digestion), as well as detoxification-related genes such as *Glutathione S-transferase 2-like*, *UDP-glucosyltransferase 2-like*, and *esterase FE4-like*. Finally, two heat shock proteins, *Tret1-like*, and *Gamma-tubulin complex component 6* are stress related, while *Toll-like receptor 6*, *tol-Pal system protein TolA-like*, *synaptic vesicle glycoprotein 2B-like*, *Histidine-rich glycoprotein-like*, and *Spaetzle* are likely immune related.

Among Ditrysia, there was a notable increase in the number of immune-related genes that provide resistance against *Bacillus thuringiensis* (Bt) crystal protein toxins. These genes include two *CD63 antigen*, *Leukocyte surface antigen CD53-like*, and *Peroxisomal membrane protein 11C-like*. Other immune related genes include *Tetraspanin-2A* and *Prostaglandin E2 receptor EP2 subtype*. Other Ditrysia-specific genes were also found to be associated with vision, chemosensation, detoxification, and stress responses, and a family, *Venom carboxylesterase-6-like,* is linked to venom production. Additionally, several genes related to mating behavior, oogenesis, and embryo development were identified, including *Zinc-type alcohol dehydrogenase-like protein SERP1785* (*ADH*), *Spherulin-2A-like*, and *Membrane-associated protein Hem*, where *ADH* is a HGT-derived gene, likely from Bacillaceae (**Figure S3**). Other HGT-derived genes in Ditrysia include *Chitinase A-like*, *Glycerophosphodiester phosphodiesterase GDPD6-like* (*GDPD*), *D-erythronate dehydrogenase-like* (*denD*), and *Cysteine synthase-like* (*CYS*) (**Figure S4-S7**). Notably, the *denD* gene tree suggests an additional HGT event from species close to *Zygaena filipendulae* (Zygaenidae) to *Achromobacter anxifer*, a potentially entomopathogenic bacterium (**Figure S6**). In contrast, none of the five Apoditrysia-specific HOGs show a clear connection to these functions.

### Validation of HGT-derived gene introduction

To further investigate the origin of the seven lineage-specific genes that are putatively derived from the horizontally transferred genes (i.e., two across Lepidoptera and five in Ditrysia), we tested whether the uniqueness of these genes resulted from gene losses in other lineages (i.e., Trichoptera and non-Ditrysia, respectively). We searched for potential pseudogenic sequences by blasting all HGT-derived genes to the genome assemblies of those species lacking these potential HGT-derived genes, using a sensitive tblastn tool. Results revealed that the few hits observed are based on low percentages of identical sequence positions (**Table S5**), suggesting that the absence of these genes are not due to recent pseudogenization. For the two putative Ditrysia-specific HGT-derived genes, *CYS* and *ADH*, whose annotated EggNOG functions are identical to some genes from non-Ditrysia species (**Table S3**), we used blast to compare the sequences of those potential orthologous genes from the non-Ditrysia species against NCBI nr database and found that those genes in non-Ditrysia are not *CYS* or *ADH* **(Table S5).** In addition, the blast result shows that only 18% of genes have hits with low percentages of identical sequence positions to reference HGT-derived genes (**Table S6**). These results suggest that genes found in non-ditrysian species annotated with the same EggNOG functions belong to different genes.

## Discussion

Our study found that many emergent gene families arose simultaneously with major evolutionary transitions in Lepidoptera. Among the hundreds of lineage-specific genes, we identified seven that were derived from horizontal gene transfer from distant organisms. These include *LigB* and *GH32*, which play important roles in plant tissue and compound digestion. Our analyses reveal that they were likely transferred from a fungus and bacteria in the ancestor of Lepidoptera, respectively. Another gene that was acquired through HGT was *CYS*, a vital gene in synthesizing cysteine and detoxifying hydrogen cyanide, an important phytotoxin in extant cyanogenic plants (13, 14). Cyanogenic glycosides – the precursors of hydrogen cyanide – are produced almost exclusively in angiosperms (41), a clade originating approximately 250-140 Ma (8). Since Ditrysia appeared around 172-137 Ma, and most species feed on angiosperms (3), the acquisition of *CYS* through horizontal gene transfer (HGT) was probably a key genomic change that helped Ditrysia specialize in angiosperms. We found that these genes, among others, closely align with host adaptations and ecological and morphological innovations, mirroring the evolutionary history of Lepidoptera as they adapted to exploit new food sources. Below, we describe some of the key takeaways we found from our results, focusing on those which we felt were most relevant.

### Host digestion and phytotoxin detoxification gene evolution: A possible key to Lepidoptera diversification

To understand how Lepidoptera diversification and their gene evolution are correlated, we examined our HOGs and identified genes showing significant repertory size change or those associated with lepidopteran clades that gained a key evolutionary innovation. We began with Trichoptera, and found that many rapidly evolving HOGs are related to detoxification, cuticle properties, and response to stress (**Table S2**). These genes likely facilitated survival in aquatic environments by supporting adaptations to factors such as temperature fluctuations and metal ion abundance (15). In Lepidoptera, we observed a consistent trend in which novel gene families associated with host plant detection and digestion – such as chemosensory and digestive genes – emerged as early as the Permian and Triassic, in Lepidoptera and Glossata, respectively. Examples include two HGT-derived genes (*LigB* and *GH32*), both associated with dietary function (see below). Given that the origin of Lepidoptera and Glossata significantly predate the angiosperm radiation – and that these genes are present in nearly all Lepidoptera species – we postulate that these genes played a role in the detection and digestion of non-angiosperm host plants. New universal gene families linked to broad ecological functions – such as circadian regulation, vision, stress responses, immunity, and venom production – were established after the origin of Glossata, approximately 233 million years ago. We hypothesize that these genes were critical in establishing ecological innovations that arose within Glossata, such as flower visitation, nectar feeding, and long-distance flight.

Newly emergent HOGs associated specifically with the origin of Ditrysia include genes related to mating behavior, oogenesis, and an expanded repertoire of detoxification and immunity genes. The emergence of gene families associated with mating behavior and oogenesis aligns with a key morphological innovation in lepidopteran evolution – the development of two distinct female reproductive openings – one for egg-laying and another for mating, a trait unique to Ditrysia (16). This adaptation likely reflects greater control over mating and oviparity, which may have facilitated specialization on specific host plants.

Defense compounds such as secondary metabolites are well known to be in much higher diversity in angiosperm species than in non-vascular plants and gymnosperms (17). Thus, the emergence of new plant detoxification gene families in Ditrysia likely explains their predominant specialization on angiosperms. For example, *CYS*, a gene known for its role in detoxifying hydrogen cyanide – a compound found in nearly all angiosperm species – was introduced to the ancestor of Ditrysia through horizontal gene transfer (see discussion below). We did not identify any new emergent HOGs associated with host plants in Apoditrysia, a result consistent with the pattern linking angiosperm feeding to the evolutionary diversification of Ditrysia.

In summary, molecular adaptations associated with dietary evolution of Lepidoptera began with a gain of gene families that help host detection and digestion, followed by the origin of specialized plant detoxification gene families after the origin of Ditrysia. Our results suggest that these molecular adaptations likely occurred before the Angiosperm Terrestrial Revolution (100-50 Ma) (9), implying that the ancestors of extant Ditrysia were genetically pre-adapted for the radiation of their host plants. In addition to host adaptation, the emergence of new gene families, such as those associated with circadian regulation, stress responses, oogenesis, and venom production influenced ancestral lepidopteran physiology, behavior, and phenology, continuously refining the life history of Lepidoptera along with diversifying host plants.

### HGT as a driver for Lepidoptera evolution

While previous studies have suggested that HGT may have played a crucial role in the ecological success of Lepidoptera (18, 19), the evolutionary origins of HGT-derived genes remain largely unknown. This is likely because HGT-derived genes are often more species-specific and restricted to younger lineages, as foreign genes typically have lower fixation rate in the recipient species and its descendant lineages (20, 21). Although repeated introductions of a neutral foreign gene might increase its chance of being fixed in a lineage, it remains unlikely to be maintained in an evolutionary ancient lineage with high diversification rate (e.g. Lepidoptera) without a significant contribution to the recipient’s fitness. In the present study, we identified HGT-derived genes that are specific to and nearly universal in Lepidoptera and Ditrysia. Since there is no evidence of gene loss in their sister lineages (**Table S5**), these HGT-derived genes were most likely introduced to their common ancestor and subsequently maintained in their descendants. These HGT-derived genes likely played a pivotal role in supporting the dietary transition from algae, diatoms, and aquatic plant debris to fungi and primitive terrestrial plants, ultimately enabling the adaptation to the most dominant angiosperm species.

Among the two HGT-derived genes that were likely introduced to the common ancestor of Lepidoptera, *LigB* appears to have horizontally transferred from a fungus that degrades aromatic compounds (22). Plant aromatic metabolites are made of many structural and volatile materials such as lignin, phenolic compounds, terpenoids, and alkaloids. Acquiring *LigB* is thus likely to function in digesting host plant tissue or detoxifying metabolic compounds. *GH32*, on the other hand, was likely introduced from a bacterium and involved in digestion, as a recent study on this gene indicates its function in breaking down host plant sucrose, fructooligosaccharides, and fructans (23). The ability to efficiently digest carbohydrates and degrade aromatic compounds supports the hypothesis that a key diet innovation took place in ancient Lepidoptera during late Carboniferous to early Permian. During this period, Lepidoptera began feeding on living fungi or plant tissues, which contained a higher proportion of simple and complex carbohydrates and presented new challenges of facing a diversity of aromatic defense compounds (24, 25).

The five HGT-derived genes introduced to the common ancestor of Ditrysia are responsible for male courtship, development, host plant digestion, and detoxification. *Zinc-type alcohol dehydrogenase-like protein SERP1785* (*ADH*, or LOC105383139) is a prevalent HGT-derived gene in Lepidoptera and plays a critical role in enhancing male courtship behavior and improving egg fertilization rate in the diamondback moth (*Plutella xylostella*) (18). This gene was likely introduced to the common ancestor of Ditrysia to aid in the control of female insemination, as ditrysian females evolved a complex reproductive system with two separate openings for mating and oviposition (26). Interestingly, *spherulin-2A-like* and *membrane-associated protein Hem*, two genes that form specific HOGs in Ditrysia, also aid in oogenesis (27–29). *Chitinase A-like* (*ChiA*) genes that encode chitinase are originally found in bacteria and aids in chitin degradation during molting (30), while *Glycerophosphodiester phosphodiesterase GDPD6-like* (*GDPD*) is associated with lipid metabolism in bacteria. In insects, *GDPD* is thought to play a role in regulating phospholipid composition during metamorphosis (31, 32). These genes were present in all ditrysian genomes that we examined but absent in non-ditrysian species. Investigating the direct functions of these two genes by manipulating their expression in larvae or pupae of different Ditrysia species could enhance our understanding of how HGT-derived genes influence the developmental process or morphology.

The putative HGT-derived gene, *D-erythronate dehydrogenase* (*denD*), found in nearly all ditrysian genomes examined, is known to synthesize vitamin B6 (pyridoxine) in bacteria (33, 34). In insects, vitamin B6 is obtained from food and symbiotic microorganisms, and it plays a key role in physiological processes, including larval growth and ovary development (35–39). The discovery that there is down-regulation of *denD* in the midgut of Lepidoptera after ingestion of phytoecdysteroids (plant-derived steroids that interfere with digestion in phytophagous insects), suggests the role of this gene in digestion (40). Finally, *CYS* is a putative HGT-derived gene that has also been identified in Diptera and Acari (e.g. dark-winged fungus gnats and mites). *CYS* plays a role in synthesizing cysteine and detoxifying hydrogen cyanide, an important phytotoxin in cyanogenic plants (13, 14). Interestingly, cyanogenic glycosides – the precursors of hydrogen cyanide – are produced almost exclusively in angiosperms (41), a clade originating approximately 250-140 million years ago (8). Given that Ditrysia originated around 172-137 Ma and most ditrysian species feed on angiosperms (3), the acquisition of *CYS* through HGT was likely a critical genomic innovation. This adaptation likely facilitated the success of Ditrysia, enabling angiosperm specialization and driving subsequent species diversification.

Nevertheless, it is noteworthy that *CYS* genes are also present in species that do not feed on cyanogenic host plants, including the highly diverse ditrysian family Tineidae, with more than 3000 described extant species and an origin of approximately 150 Ma (3). The ability to synthesize cysteine, essential for promoting caterpillar growth, may have become an important physiological innovation for Ditrysia, as cysteine is typically scarce in their diet (42). In sum, our findings reveal that five of the seven horizontally transferred genes that we identified are linked to digestion and detoxification. These genes likely played a critical role in providing adaptive advantages during the evolutionary arms race with their angiosperm hosts.

### Methodological limitations

While comparative genomics is widely recognized for providing valuable insights into the evolutionary history of living organisms, interpreting functions of poorly studied genes remains a significant challenge. Statistical approaches, such as enrichment analyses, can help identify signals present across many genes, but the absence of functional annotation for most genes remains a major hurdle, as their roles remain unknown until clarified through empirical experimentation. Another challenge is that statistical approaches can overestimate the number of potentially significant genes. In the present study, we addressed this by applying strict criteria to ensure that resulting genes were most prevalent in focal taxa and absent in others, before assessing their functional uniqueness. Although we successfully identified the origin of genes that likely played a significant role during the evolutionary history of Lepidoptera, we acknowledge that some genes may have been overlooked. While gene ontology (GO) enrichment is commonly used to infer gene function, genes lacking GO annotations might have been excluded from the analyses. We conducted an extensive search of known gene function from the published literature, focusing on studies with experimental evidence in insects. Despite these limitations, our results highlight the significance of these genes and emphasize the need for further functional validation.

## Methods

### Genomic data, gene models, and functional annotation

We collected 431 trichopteran and lepidopteran genome assemblies from NCBI GenBank and Darwin tree of life genome portal from 2022 August to 2023 November. We kept the assemblies with BUSCO completeness score higher than 90% based on the endopterygota_odb10 database (402 assemblies) using BUSCO v5.3.0 to ensure the quality of the downstream analyses (43). The high BUSCO score indicates that less missing conserved genes in the genome and thus the high integrality of the assemblies is advised. For those genome assemblies with more than 90% complete endopterygota BUSCO genes, we reran BUSCO with lepidoptera_odb10 database to investigate the evolution of single copy genes along the evolutionary history of Amphiesmenoptera (Trichoptera+Lepidoptera). The complete scores are aggregated by the superfamily and plotted with species count of the superfamily extracted from previous estimates (10, 44) (**Figure 1**). For genome re-annotation, we selected 62 representative species from 6 Trichoptera and 21 Lepidoptera superfamilies (from 6 and 42 families, respectively) with the highest BUSCO score in the family (**Table S1**). For each genome, we built gene models using BRAKER3 EP (proteins) or ETP (proteins + RNA-Seq) pipeline depending on the availability of the transcriptome RNA data in the NCBI SRA database (**Table S1**) (45–56). The primary transcripts of the AUGUSTUS gene models (selected by “primary_transcript.py”) from BRAKER3 were used to perform functional annotation using EggNOG v 2.1.6 with HMM search against EggNOG Arthropoda database (eggnog_5.0, Arthropoda 6656) and DIAMOND v2.0.9 function blastp against RefSeq non-redundant protein database with e-value cutoff of 1×10^-10^ (57).

### Phylogeny inference

We reconstruct phylogeny with the 62 recruited species using lepidoptera single copy orthologs inferred from BUSCO. Specifically, we collected single copy orthologs for each species and identified 1,439 genes presenting in all the 62 species. For each gene, we align the sequences using MAFFT v7.490 with L-INS-i aligning strategy to generate multiple sequence alignments and subsequently concatenate the alignments into a supermatrix alignment (58). We employed IQ-tree v2.0.3 to infer Maximum Likelihood (ML) tree based on the supermatrix alignment with each gene being partition unit for best substitution model selection using PartitionFinder (59, 60). We then assessed branch support for the ML tree using 1,000 replicates SH-aLRT support and ultrafast bootstrap (61). Finally, we converted the species tree into ultrametric tree using treePL, with the divergence time calibrations of crown ages of Amphiesmenoptera, Lepidoptera, Trichoptera, Ditrysia, and Apoditrysia estimated in the recent Lepidoptera phylogenomic analysis (3, 62).

### Orthogroup inference and gene content evolution

We use the primary transcript of predicted genes of 62 species to perform orthologs inference using OrthoFinder v2.5.2 (63). We choose phylogenetic hierarchical orthogroups (HOGs) of all species (N0.tsv) for downstream gene family evolution analyses. For each HOG, we checked the absence and presence of genes from species in the taxa of Trichoptera, Lepidoptera, Glossata, Ditrysia, and Apoditrysia. Lineage-specific HOGs were identified as being present in >95% species of focal taxa and its nested taxa, and completely absent in other species. Rapid gene repertoire size changes (i.e., expansion and contraction) of focal taxa were assessed using CAFE v5.0.0 with the use of ultrametric species tree and inferred HOG gene counting matrix (64, 65). Two gamma categories and a significant threshold of p ≤ 0.01 were applied to the analysis.

### Functional annotation of HOGs

To obtain the putative function of HOGs, we annotated all HOGs with consensus EggNOG functions and the best blast results of all the genes (majority rule, and duplicated genes were included) in the HOG. Since many HOGs are annotated with the same EggNOG function but may not have identical annotation convention (e.g. P450 vs. p-450), we clustered functions by dividing the functional annotations (the description of the function) into individual words which were separated by space and non-alphanumeric characters and used these words to match other annotations to allow words being reordered and case insensitive. The annotations in the same keyword searching cluster are treated as the same putative function. To summarize the functions of these lineage specific HOGs, we extracted GO terms of all the genes in each HOG from EggNOG annotations, and selected GO terms that presented in genes from more than 80% of the species in the group to remove the inconsistent annotations. The selected GO terms were used to further summarize their biological process functions using Revigo with an assigned database from *Drosophila melanogaster* (66). We selected a small resulting size and kept the terms with frequency less than 10% to focus on the more specific terms (**Supplemental Data 1**). Finally, we manually annotated the putative function of lineage specific genes by searching their blast and EggNOG function annotation across published literatures and the function was determined by experimental tests such as gene expression on insects (**Table 1**).

**Table 1.** Lineage-specific gene families in three major groups of Lepidoptera exhibit known or putative functions associated with key ecological traits. These gene families reflect adaptations to their respective environments and life strategies, including host plant specialization, stress response, and morphological innovations, among others.

| Putative functions | Lepidoptera (276 Ma) | Glossata (233 Ma) | Ditrysia (157 Ma) |
| --- | --- | --- | --- |
| Chemosensory | chemosensory protein (Insect pheromone-binding family, A10/OS-D) | glutamate receptor 1-like | ejaculatory bulb-specific protein 3-like (Insect pheromone-binding family, A10/OS-D) |
|  |  | pickpocket protein 28-like | steroid hormone receptor ERR1 |
|  |  | odorant receptor | ionotropic receptor 75a-like |
|  |  | homeotic protein empty spiracles-like |  |
| Digestion | Catalytic LigB subunit of aromatic ring-opening dioxygenase | unnamed protein product with peptidase inhibitor activity | unnamed protein (Trypsin-like serine protease) |
|  | sucrose-6-phosphate hydrolase-like | trypsin-7 isoform X1 | serine protease gd-like |
|  | serine/threonine-protein kinase unc-51 | proton-coupled folate transporter-like | D-erythronate dehydrogenase-like |
|  | membrane alanyl aminopeptidase-like | Myrosinase 1 |  |
| Detoxification | cytochrome P450 4C1-like | esterase FE4-like | cysteine synthase-like |
|  | unnamed protein product (Carboxylesterase family) | UDP-glucosyltransferase 2-like | unnamed protein (Sulfotransferase family) |
|  | Flavin containing amine oxidoreductase | glutathione S-transferase 2-like | probable cytochrome P450 49a1 |
| Immunity | toll-like receptor 4 | unnamed protein product (Spatzle) | prostaglandin E2 receptor EP2 subtype |
|  |  | trichohyalin-like | tetraspanin-2A |
|  |  | histidine-rich glycoprotein-like | CD63 antigen |
|  |  | toll-like receptor 6 | CD63 antigen-like isoform X2 |
|  |  | tol-Pal system protein TolA-like | leukocyte surface antigen CD53-like |
|  |  | NADPH oxidase 4-like | peroxisomal membrane protein 11C-like |
| Cuticle property | Chitin-binding domain type 2 | peroxisomal N(1)-acetyl-spermine/spermidine oxidase-like |  |
|  | pyrimidodiazepine synthase-like | Endocuticle structural glycoprotein ABD-5-like |  |
|  |  | Endocuticle structural glycoprotein ABD-4-like |  |
| Response to stress | Lanthionine synthetase C-like | gamma-tubulin complex component 6 | STE20/SPS1-related proline-alanine-rich protein kinase |
|  |  | alpha-crystallin A chain-like | facilitated trehalose transporter Tret1-like |
|  |  | heat shock protein 22-like |  |
| Mating |  |  | zinc-type alcohol dehydrogenase-like protein SERP1785 |
| Oogenesis |  |  | spherulin-2A-like |
|  |  |  | membrane-associated protein Hem |
| Circadian |  | circadian clock-controlled protein daywake-like |  |
| Vision |  | opsin-1 | retinol dehydrogenase 11-like |
|  |  | disco-interacting protein 2 isoform X2 |  |
| Venom component |  | venom dipeptidyl peptidase 4 | venom carboxylesterase-6-like |

### Source and prevalence of lineage specific HOGs

We further investigated the potential sources of these new emergent, lineage specific HOGs in the focal group. First, we counted HOGs that were annotated with the same function (defined above) in all species, focal taxa, and their sister counterpart. This provides information about how prevalent and diverse the functions are. The lineage specific HOGs with more unique function (with function less commonly seen in other HOGs) implies its importance to acquire the new emergent ortholog. Vice versa, the lineage specific HOGs that share common function with many other HOGs are more likely to be resulted from the frequent gene duplications and losses. We used Fisher’s exact test to quantify the functional uniqueness of the lineage specific HOGs by testing if the odds ratio is significantly less than one using the “fisher.test” function in R (67), where the odds ratios were calculated in equation 1. The lower *p*-value indicates that the focal taxon has an exceeding number of new emerging HOGs for the function.

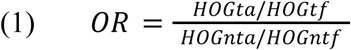

In equation 1, *OR* is the odds ratio of the focal HOG; *HOGta* is the number of HOGs annotated with target function from all species; *HOGtf* is the number of HOGs annotated with target function only found in the focal taxon; *HOGnta* is the number of HOGs annotated with non-target function from all species; and *HOGntf* is the number of HOGs annotated with non-target function only found in the focal taxon.

For the putative HGT-derived genes, we tested if the lineage specific genes were acquired from HGT by blasting all the genes in the lineage specific HOGs to the genes in the lepidopteran HGT database with an e-value cutoff of 1×10^-10^ (18). For those lineage specific HOGs that contain all their genes blasting to HGT-derived genes, we further test if the genes were indeed acquired during their common ancestors, or alternatively, the result of gene loss in the other lineages. Specifically, we searched for the potential psedogenized sequences in the genomes of those species without HGT-derived genes. This was done by blasting the HGT protein sequences to the genomes of those species without HGT using more sensitive tblastn which can detect disconnected and possibly defunction coding sequences in the genome (68). Results of multiple hits in the genome with high percentages of identical positions implies recent gene loss in the species and thus the time of HGT could have occurred earlier than the common ancestor of the lineage. In addition, since there are two HGT-derived genes in Ditrysia, *CYS* and *ADH*, having their annotated EggNOG functions identical to other genes from non-Ditrysia species (see results), we further tested if those genes from the non-Ditrysia species are also from HGT but were placed in different HOGs and thus have not tested for HGT. For this task we blast genes from these HOGs against the lepidopteran HGT database and the NCBI nr database for protein product annotation using DIAMOND v2.0.9 blastp (51). For detecting the potential gene donators, we further blast all the HGT-derived gene sequences against NCBI nr database using DIAMOND v2.0.9 blastp but excluding hits from Lepidoptera species. The best 5 hits of each sequence (if any) were collected and used to create sequence alignments after removing duplicated hits using MAFFT v7.490 (58). Gene trees of the seven HGT-derived genes were reconstructed using IQ-tree v2.0.3 with the best substitution model selected based on PartitionFinder embedded in IQ-tree v2.0.3 (59, 60). The branch supports were assessed using SH-aLRT support and ultrafast bootstrap with 1,000 replicates (61).

## Acknowledgements

The authors thank UFIT Research Computing for providing computational resources and support that contributed to the research results presented in this publication. We also thank Dr. R. Keating Godfrey for the valuable advice and guidance throughout the research. For more information, visit http://www.rc.ufl.edu. Funding for this project was provided by National Science Foundation (NSF) DEB #2624250, EF #2217159, and IOS #1920895.

