## Supplemental FigureS1-S7 for "Gene family evolution reveals dietary adaptations in butterflies and moths"

Paul B. Frandsen: 0000-0002-4801-7579

Akito Y. Kawahara: 0000-0002-3724-4610

Running Title: Evolutionary Genomics of Lepidoptera

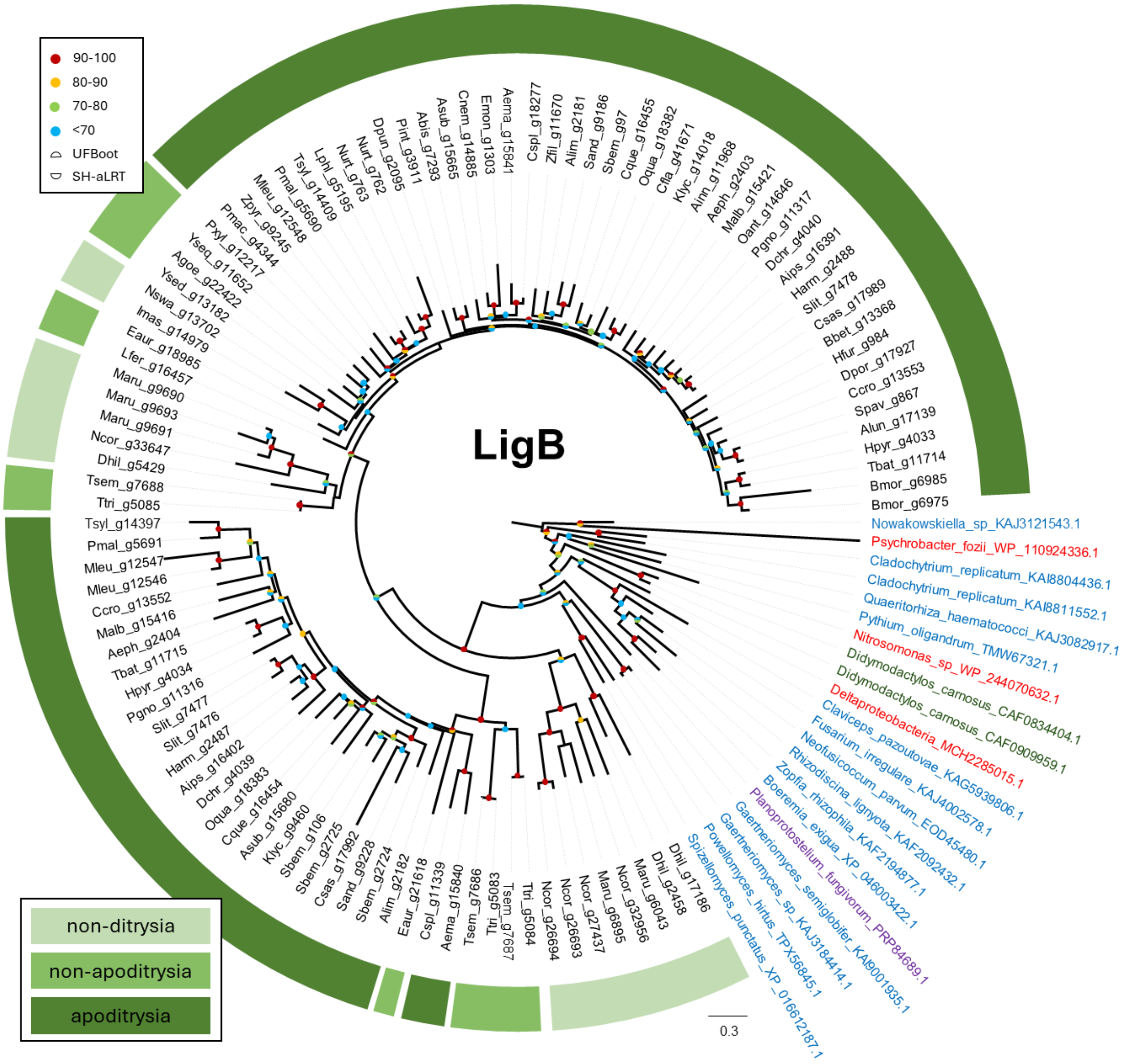


**Figure S1**. *Catalytic LigB subunit of aromatic ring-opening dioxygenase* (*LigB*) gene tree. The outgroup from the top 5 hits from the NCBI nr database includes fungi (blue), bacteria (red), Mycetozoa (purple), and Spiralia (*Didymodactylos carnosus*, green). The gene seems to undergo duplication soon after being transferred to Lepidoptera, as most two major clades have similar arrangement.

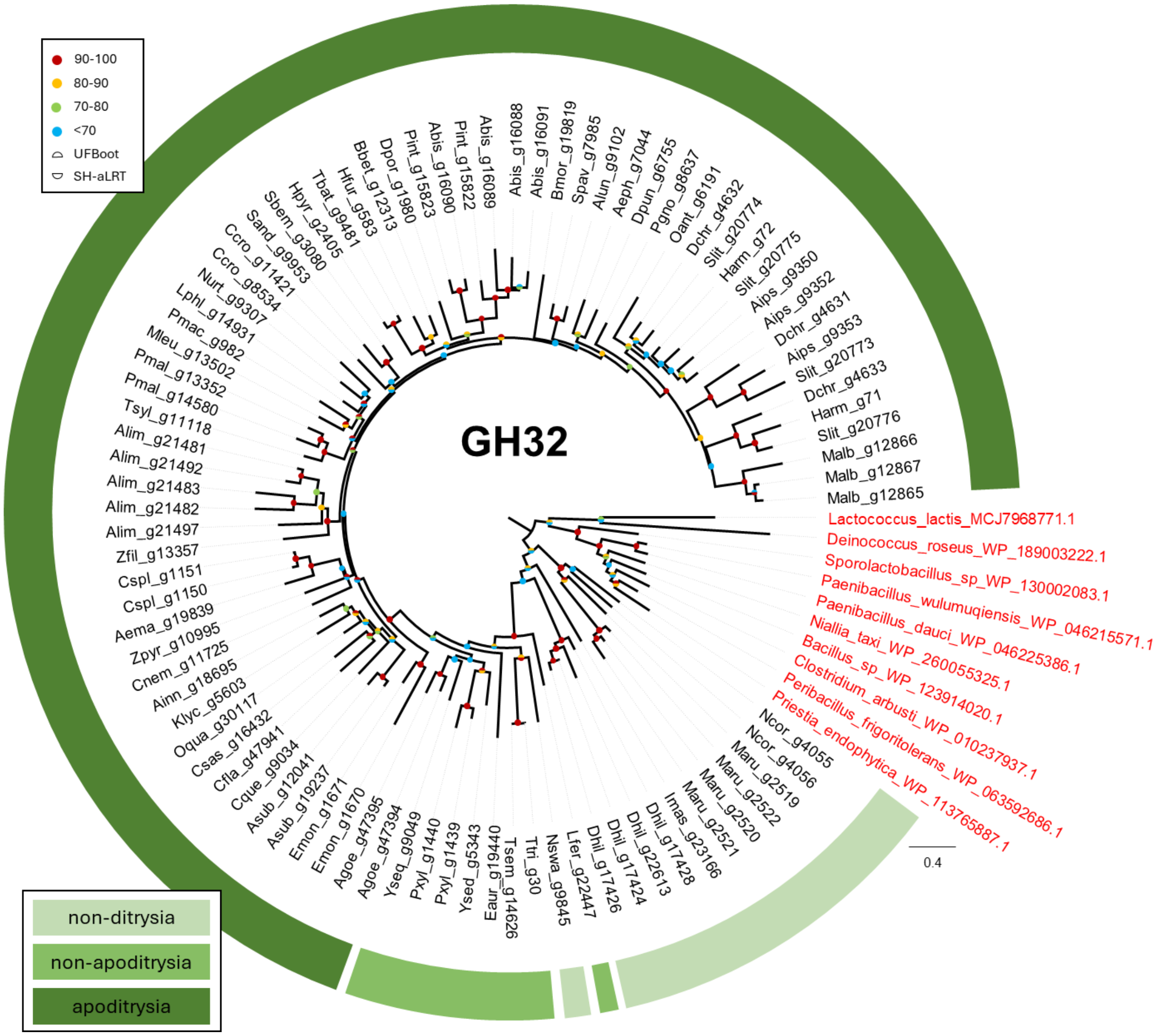


**Figure S2**. *Glycosyl hydrolases family 32* (*GH32*) gene tree. The outgroup from the top 5 hits from the NCBI nr database are all bacteria (red). The orthologs of this gene are found in both Ditrysia and non-Ditrysia species.

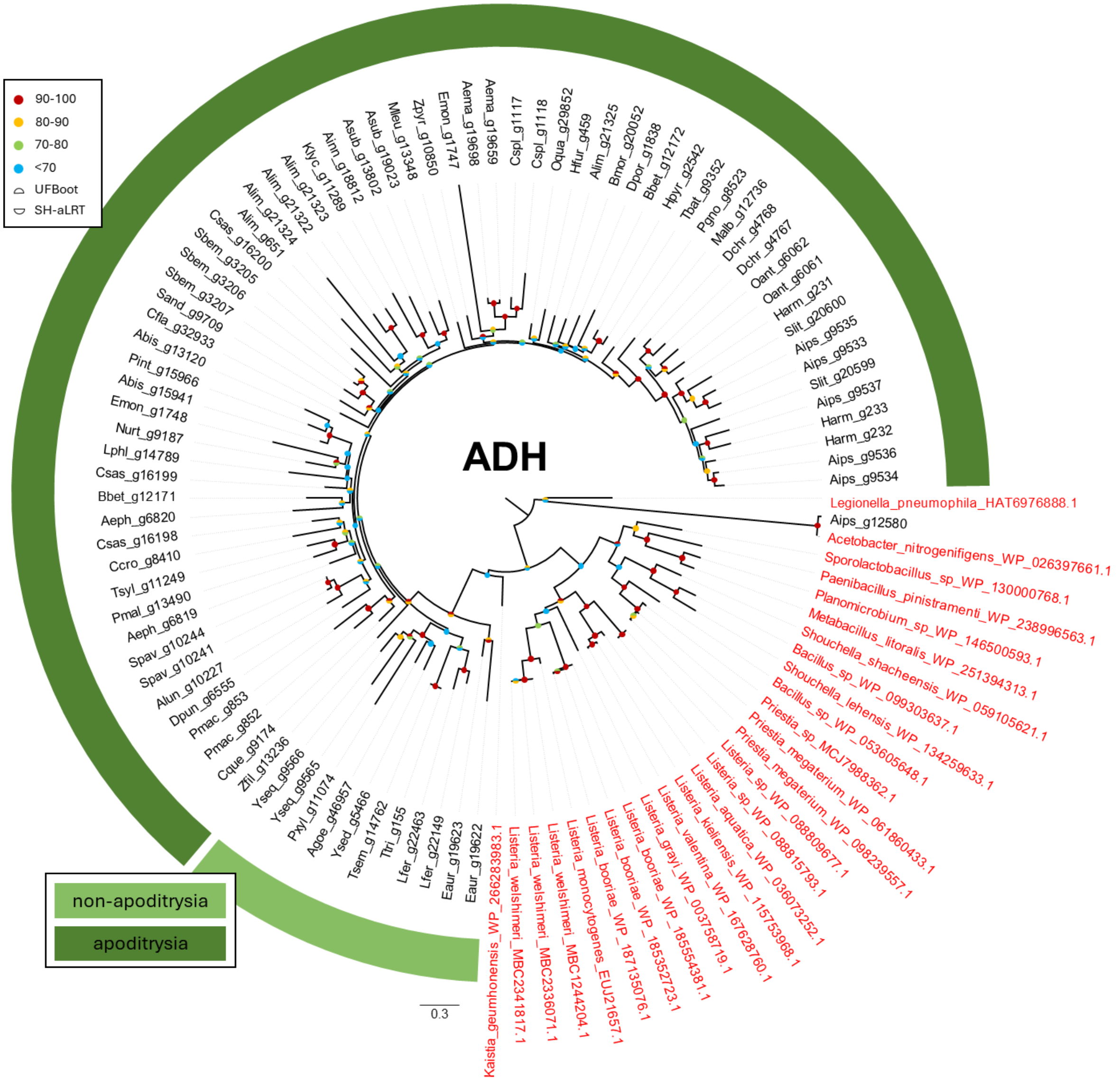


**Figure S3**. *Zinc-type alcohol dehydrogenase-like protein SERP1785* (*ADH*) gene tree. The outgroup from the top 5 hits from the NCBI nr database are all bacteria (red), with one exception that seems to be from an independent introduction from *Legionella* to *Agrotis ipsilon* (Aips_g12580).

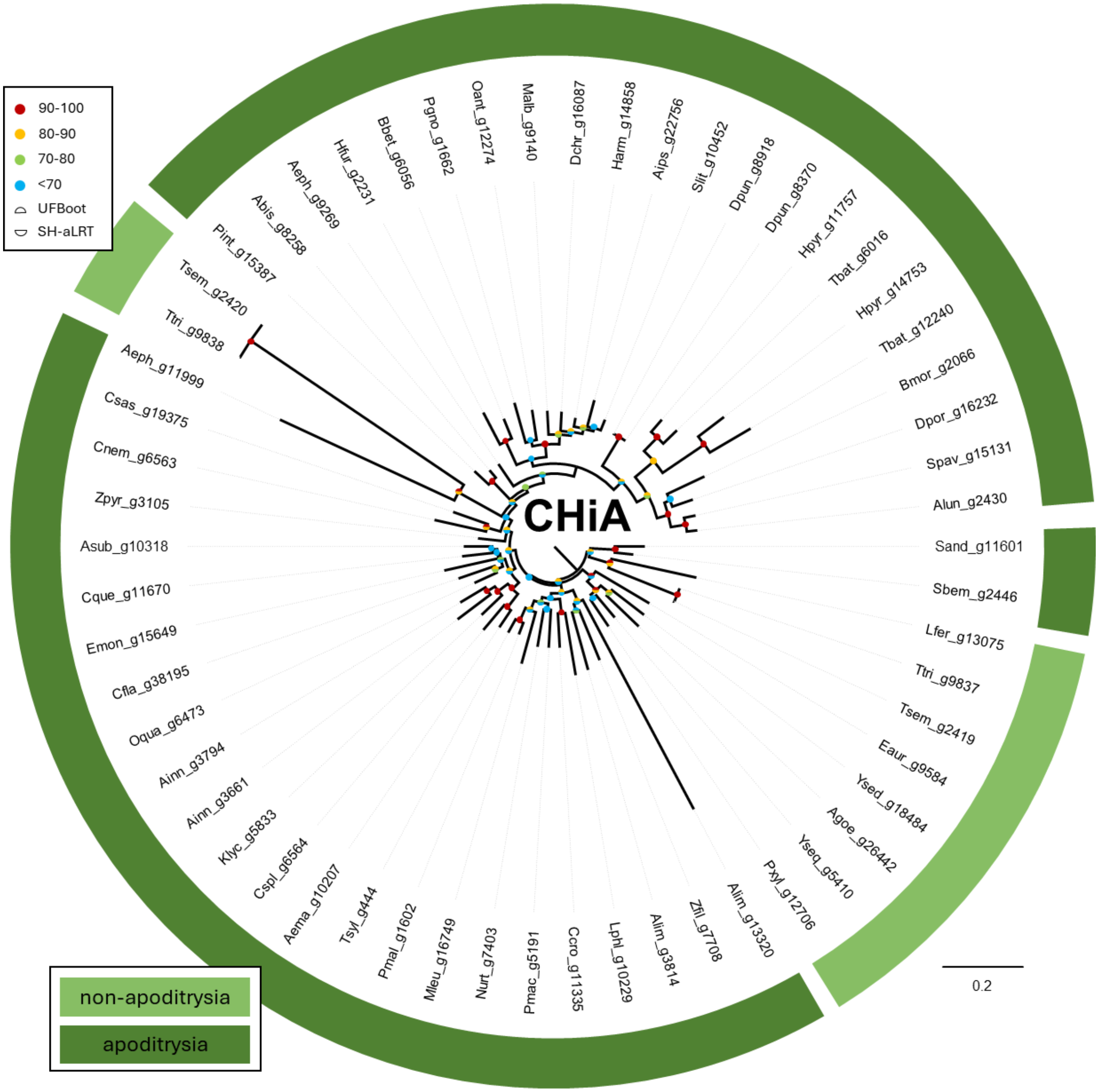


**Figure S4**. *Chitinase A-like genes* (*ChiA*) gene tree. There is no hit from the blast result thus the potential gene donator is unknown.

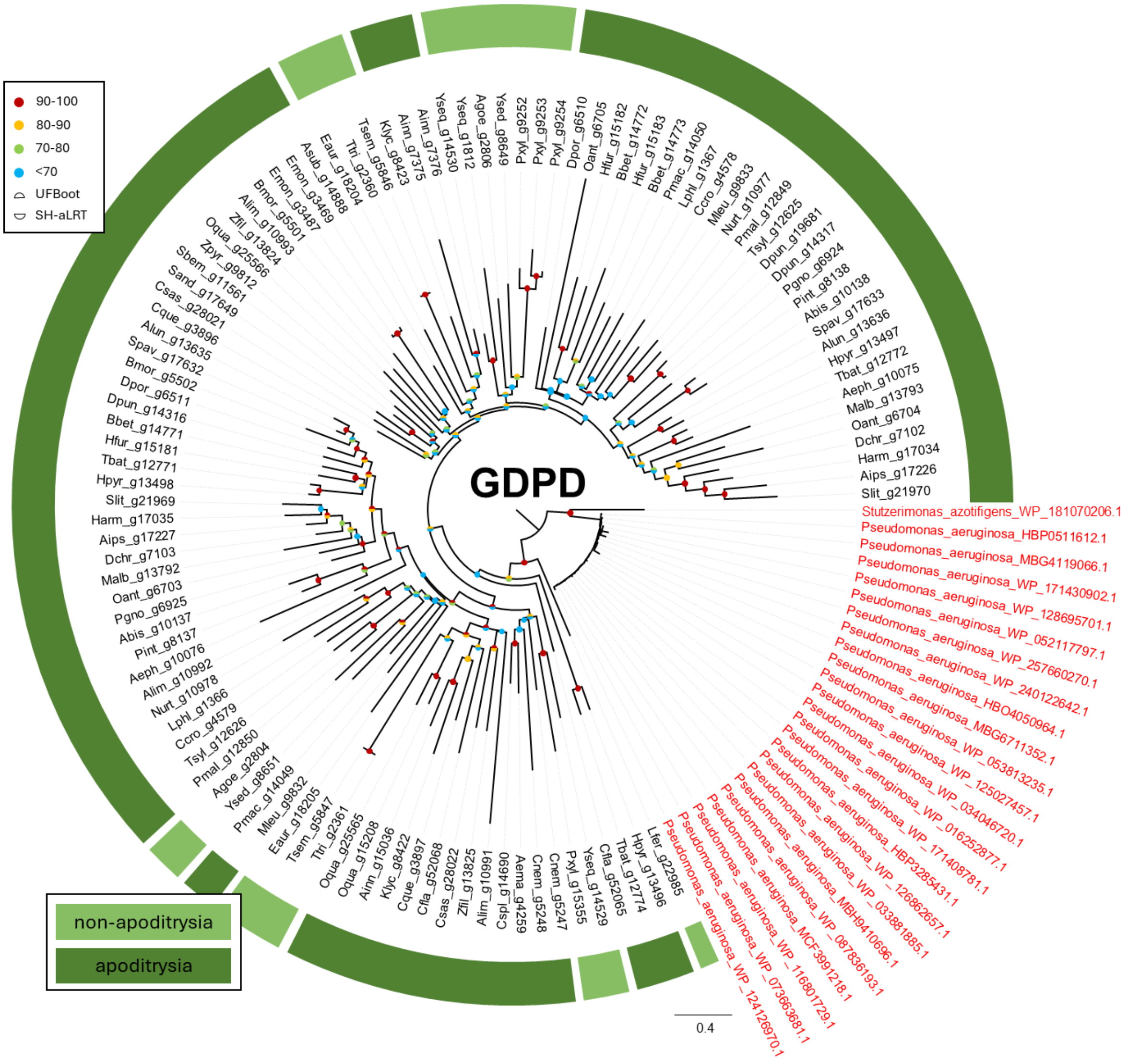


**Figure S5**. *Glycerophosphodiester phosphodiesterase GDPD6-like* (*GDPD*) gene tree. The outgroup from the top 5 hits from the NCBI nr database are all bacteria (red), mostly from *Pseudomonas* spp. Several duplications have occurred in Lepidoptera lineages.

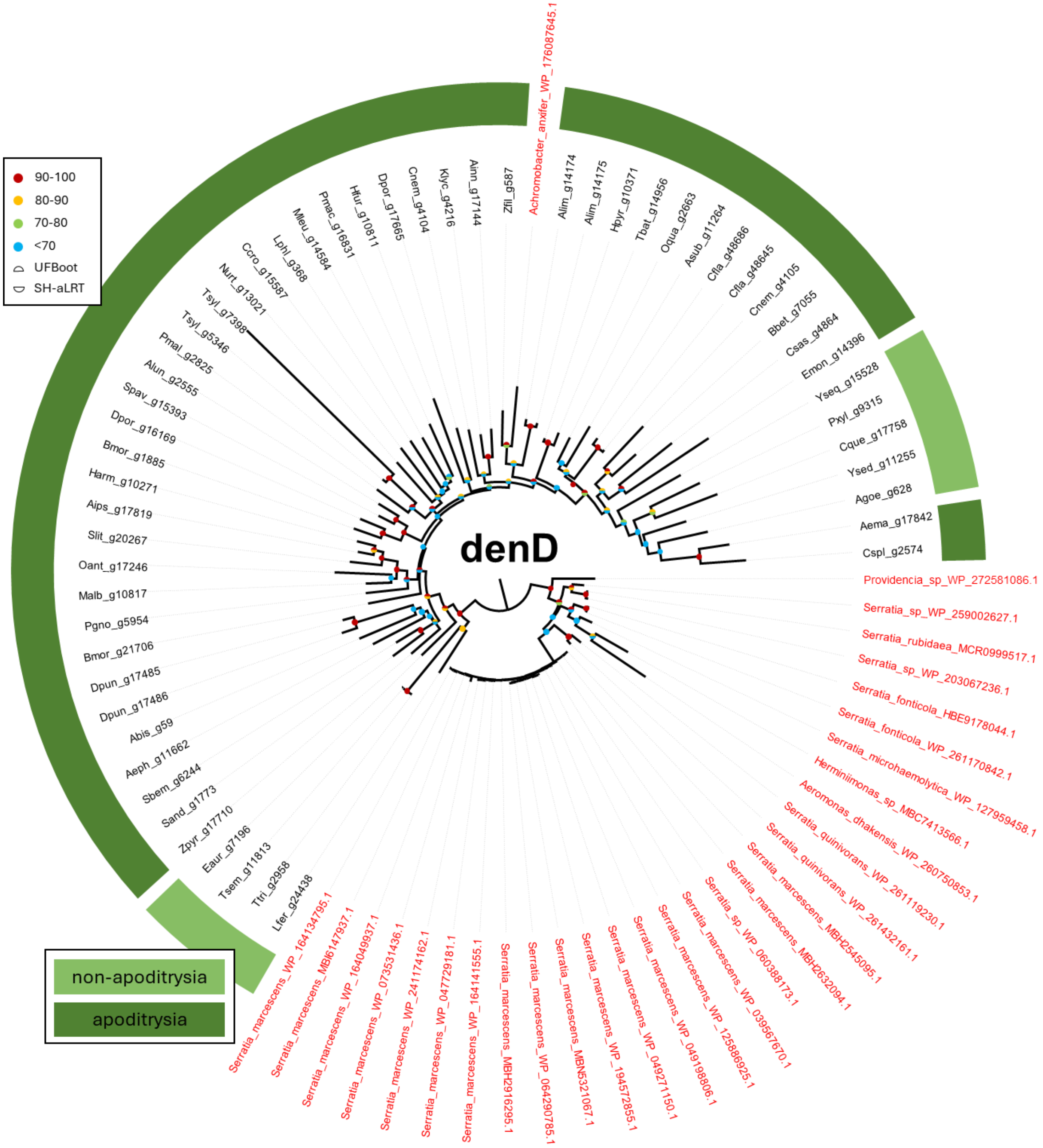


**Figure S6**. *D-erythronate dehydrogenase* (*denD*) gene tree. The outgroup from the top 5 hits from the NCBI nr database are all bacteria (red), mostly from *Serratia* spp. There seems to be an additional HGT event from species close to *Zygaena filipendulae* to *Achromobacter anxifer*.

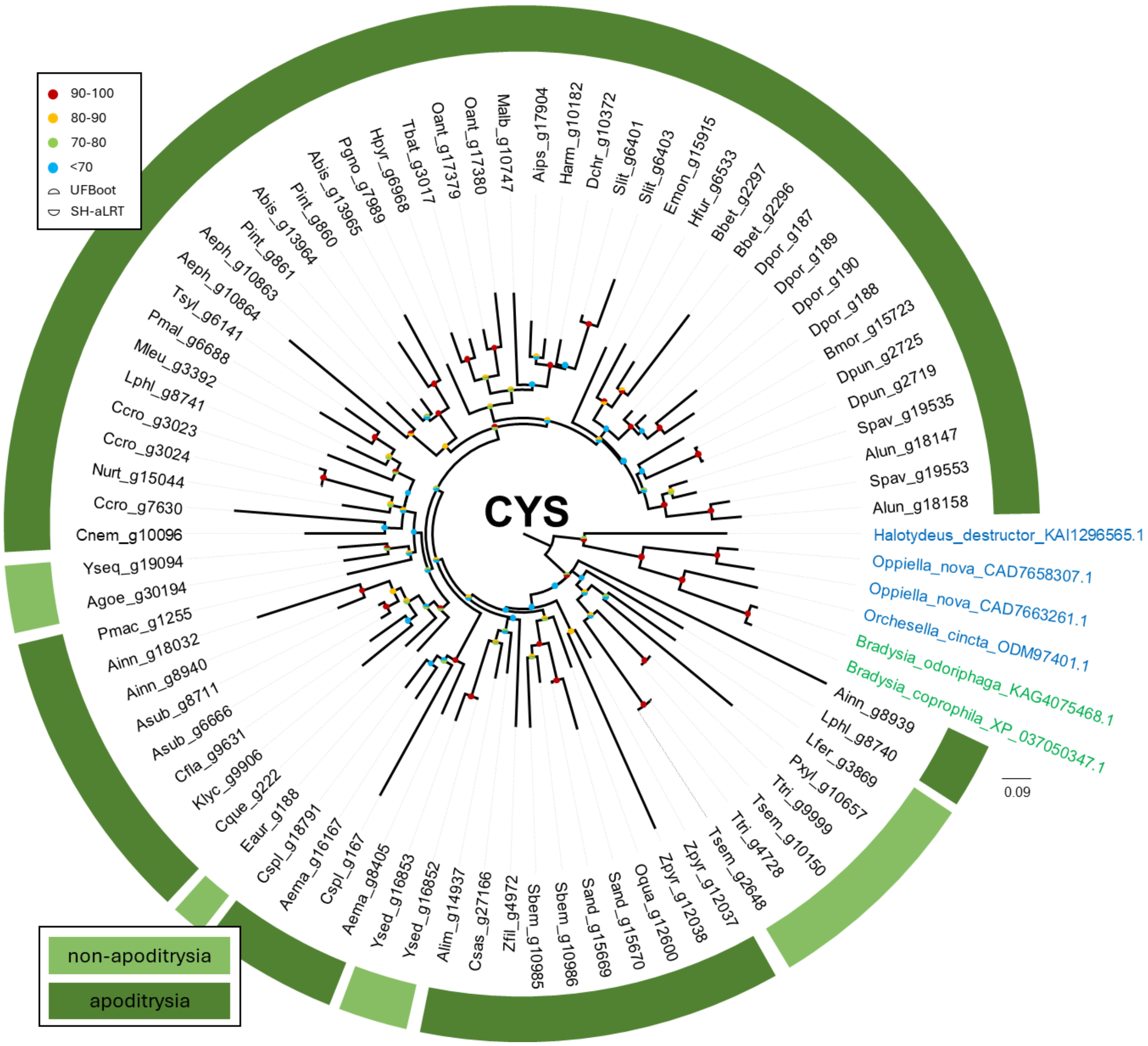


**Figure S7**. *Synthesizing cysteine* (*CYS*) gene tree. The orthologs of the gene are found only in Ditrysia. The outgroup from the top 5 hits from the NCBI nr database are fungus gnat *Bradysia coprophila* (green) and mites *Halotydeus* and Oppiella (blue).
